# *S. cerevisiae* Srs2 helicase ensures normal recombination intermediate metabolism during meiosis and prevents accumulation of Rad51 aggregates

**DOI:** 10.1101/524413

**Authors:** Laura J Hunt, Emad Ahmed, Hardeep Kaur, Jasvinder Ahuja, Lydia Hulme, Ta-Chung Chou, Michael Lichten, Alastair SH Goldman

## Abstract

We investigated the meiotic role of Srs2, a multi-functional DNA helicase/translocase that destabilizes Rad51-DNA filaments, and is thought to regulate strand invasion and prevent hyper-recombination during the mitotic cell cycle. We find that Srs2 activity is required for normal meiotic progression and spore viability. A significant fraction of *srs2* mutant cells progress through both meiotic divisions without separating the bulk of their chromatin, although sister centromeres often separate. Undivided nuclei contain aggregates of Rad51 colocalized with the ssDNA-binding protein RPA, suggesting the presence of persistent single-strand DNA. Rad51 aggregate formation requires Spo11-induced DSBs, Rad51 strand-invasion activity, and progression past the pachytene stage of meiosis, but not the DSB end-resection or the bias towards inter-homologue strand invasion characteristic of normal meiosis. *srs2* mutants also display altered meiotic recombination intermediate metabolism, revealed by defects in the formation of stable joint molecules. We suggest that Srs2, by limiting Rad51 accumulation on DNA, prevents the formation of aberrant recombination intermediates that otherwise would persist and interfere with normal chromosome segregation and nuclear division.

## Introduction

The production of haploid gametes during meiosis requires the ordered segregation of the diploid genome, such that each chromosome is present as a single copy in daughter cells. This occurs through two successive nuclear divisions (meiosis I and meiosis II) that follow a single round of DNA replication. In most organisms, homologous chromosomes of different parental origin (called homologues), each comprising two sister chromatids held together by cohesin, are first paired and then synapsed end-to-end by a structure called the synaptonemal complex (SC; reviewed in Heyting, 1996). Homologues become covalently linked by crossovers (COs), and this linkage allows their correct orientation on the meiosis I spindle, so they segregate to opposite poles during the first meiotic nuclear division. Sister chromatids are subsequently separated during the meiosis II. In most eukaryotes, homologous recombination (HR) is critical during meiosis, both for the pairing/synapsis processes that bring homologues together, and for the formation of crossovers that allow their correct segregation (reviewed in Petronczki et al., 2003).

Meiotic HR is initiated by DNA double-strand breaks (DSBs), formed by the Spo11 protein (Keeney et al., 1997). DSBs are resected, in a process dependent on Sae2/Com1 (McKee and Kleckner, 1997; Prinz et al., 1997) to form 3’ single-stranded DNA (ssDNA) capable of a homology search. During the mitotic cell cycle, HR requires the strand exchange protein Rad51, which displaces the ssDNA-binding RPA protein complex and mediates strand invasion into homologous duplex DNA (reviewed in Morrical, 2015). The invading single strand displaces a strand of the homologous duplex and primes DNA synthesis to form a displacement loop (D-loop). D-loop collapse, followed by reannealing to ssDNA at the other break end forms exclusively non-crossover (NCO) products in a process called synthesis-dependent strand annealing (SDSA; Nassif et al., 1994). Alternatively, capture of the second ssDNA by the D-loop forms a double Holliday junction (dHJ) intermediate, which can be resolved either as an NCO or as a CO (Szostak et al., 1983). During the mitotic cell cycle, most Rad51-mediated HR occurs between sister chromatids (Bzymek et al., 2010; Kadyk and Hartwell, 1992), and events that involve homologues produce NCOs, rather than COs (Ira et al., 2003; Lichten and Haber, 1989), thus reducing the risk of genome rearrangement and loss of heterozygosity.

In contrast, interhomologue COs are an important outcome of HR during meiosis, since they mediate proper homologue orientation and segregation. In many organisms, including budding yeast, meiosis-specific modifications of recombination both encourage strand invasion between homologues and production of a higher proportion of COs. One of these involves the expression of a second strand exchange protein, Dmc1 (Bishop et al., 1992). Rad51 is still present in meiotic cells, and is an essential cofactor for normal Dmc1 loading and strand invasion activity (Cloud et al., 2012). However, in budding yeast, Rad51 strand exchange activity is dispensable for recombination (Cloud et al., 2012), and in fact is negatively regulated during meiosis by at least two known mechanisms involving the meiosis-specific kinase Mek1. Mek1 kinase is activated by binding to a phosphorylated form of the meiotic chromosome axis protein Hop1 (Niu et al., 2005), which is formed in the vicinity of meiotic DSBs by DNA damage response kinases (Carballo et al., 2008). Active Mek1 phosphorylates Rad54, a protein required for Rad51 to form stable strand invasion products, and prevents Rad54-Rad51 interaction (Niu et al., 2009). Mek1 also phosphorylates and stabilizes the meiosis-specific Hed1 protein, which binds to Rad54 and prevents it from interacting with Rad51 (Callender et al., 2016).

Rad51 is also negatively regulated by the budding yeast Srs2 protein, a 3’ to 5’ DNA helicase/translocase of the UvrD family (reviewed in Niu and Klein, 2017). Srs2 interacts with Rad51 filaments *in vitro* and strips them from DNA (Kaniecki et al., 2017; Krejci et al., 2003; Veaute et al., 2003). Loss of Srs2 activity in mitotic cells leads to DNA damage sensitivity, genome instability, reduced DSB repair efficiency, and an increase in COs among repair products (Elango et al., 2017; Ira et al., 2003; Lawrence and Christensen, 1979; Marini and Krejci, 2010; Rong et al., 1991). As *srs2* sensitivity to DNA damage is partially suppressed by deletion of *RAD51*, it is thought that *srs2* mutant phenotypes relate to failures in Rad51 removal from ssDNA (Ira et al., 2003; Krejci et al., 2003). Srs2 also unwinds branched DNA structures *in vitro*, including those mimicking D-loop recombination intermediates, consistent with a role in promoting SDSA (Dupaigne et al., 2008; Kaniecki et al., 2017; Liu et al., 2017; Marini and Krejci, 2012). However, this function has yet to be fully investigated *in vivo*. In meiosis, Srs2 activity is required for normal spore viability and meiotic progression, and *srs2*Δ mutants show reduced formation of COs and NCOs (Palladino and Klein, 1992; Sasanuma et al., 2013a; Sasanuma et al., 2013b).

We have analysed further the importance of Srs2 function during meiosis and found that it is required for normal recombination intermediate metabolism and nuclear division. In *srs2* mutants, Rad51 protein appears in aggregates after exit from pachytene, when the SC has been dissolved. These Rad51 aggregates are often associated with RPA, and arise only if programmed DSBs are formed and if Rad51 has full strand exchange capability. *srs2* mutants show partial defects in meiotic nuclear divisions, but cytological investigation of chromosomal segregation implies that sister centromere separation occurs normally. These data suggest that *srs2* mutants suffer entanglements caused by abnormal interhomologue recombination intermediates. Consistent with this, we found evidence for defects in formation of stable interhomologue recombination intermediates. We propose that loss of Srs2-mediated negative regulation of Rad51 allows for defects in DNA interactions during pachytene, which later lead to defects that prevent normal nuclear division.

## Results

### Sporulation is delayed and reduced in srs2 mutants, with decreased spore viability

Known meiotic defects caused by loss of Srs2 activity include nuclear division defects, reduced sporulation and reduced spore viability (Palladino and Klein, 1992). We confirmed these phenotypes in our *srs2* mutant strains, which included: *srs2*Δ, which produces no Srs2 protein; *srs2-101*, which produces a protein lacking ATP hydrolysis activity (Rong et al., 1991); and *srs2-md*, a replacement of the endogenous *SRS2* promoter with the *CLB2* promoter, where *SRS2* is expressed in mitotic cells but not during meiosis (Chu et al., 1998).

All three *srs2* mutants showed meiotic nuclear division defects. The fraction of cells completing both meiotic nuclear divisions was reduced from 94% in *SRS2* strains to 60%, 59% and 60%, respectively in cells homozygous for *srs2-md*, *srs2-101* and *srs2*Δ. In addition, nuclear division was delayed by approximately 1 h, and a substantial fraction (~20%) of *srs2* mutant cells completed only one of the two nuclear divisions (Fig. 1a-e). Incomplete nuclear division could also be observed in *srs2* strains, with nuclei remaining connected by chromosome bridges at times when nuclear division was complete in wild type (Fig. 1f). Spore viability is reduced in cells that complete meiosis, from 98% in wild-type to 63-68% in *srs2* mutants (Fig. 1g). Tetrad spore death patterns typical of defects specific to either meiosis I or meiosis II were not observed, suggesting that segregation failure is not specific to a single stage of meiosis (Fig. 1h).

**Fig. 1.**
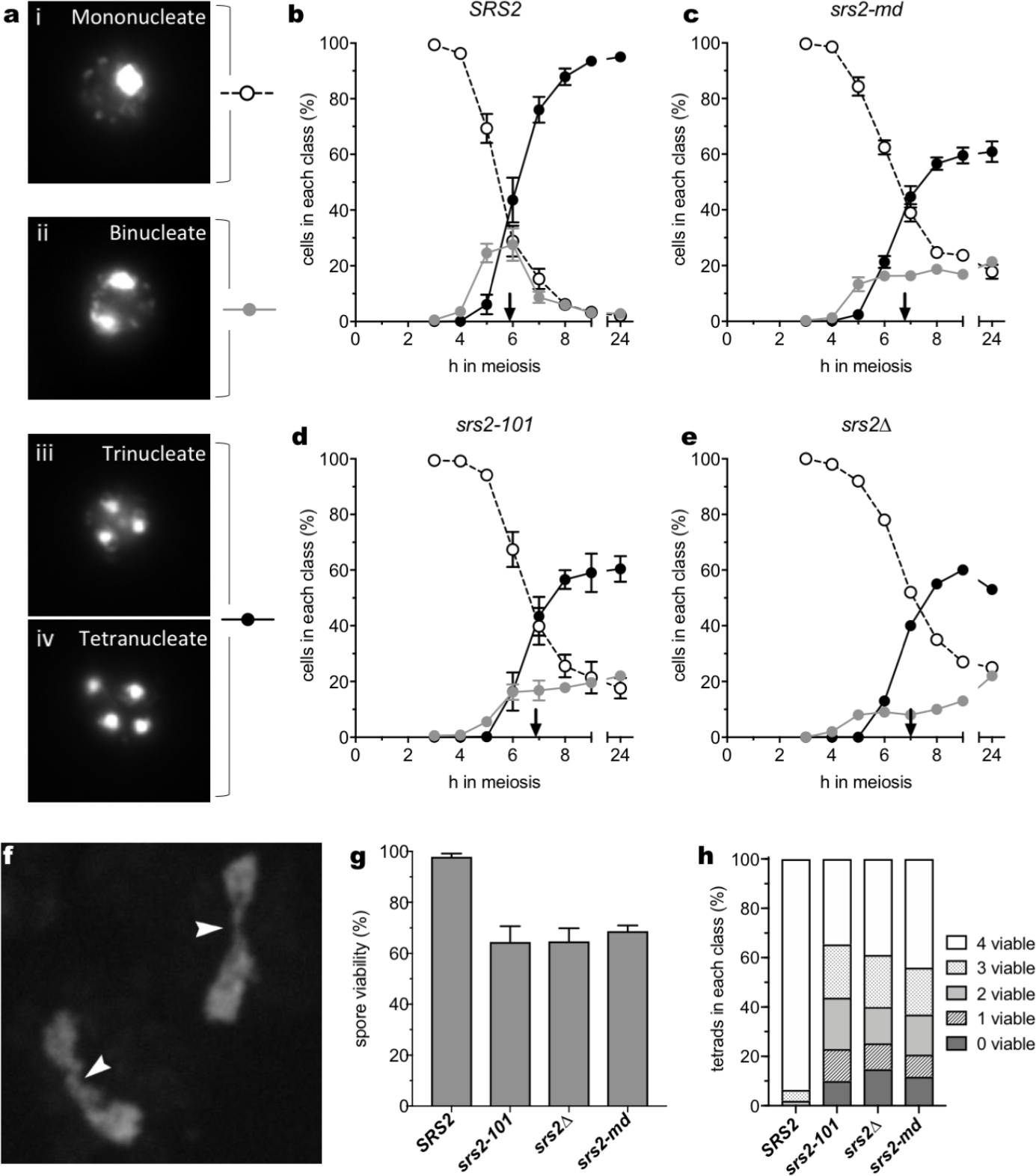
Sporulation defects caused by loss of Srs2 activity. (a) Examples of nuclear morphology. (b) In *SRS2* strains, about 95% of cells progress through the two meiotic nuclear divisions. (c-e) Progression through nuclear divisions is reduced to about 60% compared to wild-type and is delayed by ~ 1h; vertical arrows on X-axis indicate when 40% of cells had completed meiosis II (*SRS2*, n=7; *srs2-md*, n=7; *srs2-101*, n=5; *srs2*Δ, n=1; error bars— standard error of the mean). *srs2* mutants also display persistent binucleate cells. (f) DNA bridges observed in in *srs2* mutants (example from 9h, *srs2-101*). (g) Reduced spore viability in *srs2* mutants. (h) Patterns of spore viability in tetrads from *srs2* mutants are inconsistent with either meiosis I or meiosis II nondisjunction. Numbers of independent experiments and total tetrads analysed are: *SRS2* - 5, 159; *srs2-101* - 2, 99; *srs2Δ* - 2, 100; *srs2-md* - 3, 139. Error bars in panel g indicate range.

### The spindle cycle continues in srs2 mutants despite nuclear division defects

During each round of meiosis, the spindle pole body (SPB, budding yeast centrosome equivalent) must duplicate, divide and migrate to opposite poles of the cell in order for the dividing chromosomes to be correctly drawn along the tubulin spindles at anaphase. Duplicated SPBs are initially connected by a bridge (Byers and Goetsch, 1975), and spindle separation is controlled by activity of the cyclin-dependent kinase Cdc28/Cdk1 (Jaspersen et al., 2004). During the meiosis I to meiosis II transition, SPBs must be relicensed for duplication in a process regulated by the Cdc14 phosphatase (Fox et al., 2017). Thus, analysis of SPB division provides a useful indicator of meiotic cell cycle progression.

To monitor SPB division, we examined spread cells from strains expressing fluorescently tagged SPB (Cnm67-mCherr*y*) and spindle proteins (GFP-Tub1), neither of which altered meiotic progression (Fig. S2b). Cells were classified by number of SPB signals, as for nuclear division (Fig. 2a). As expected, due to SPB division preceding nuclear division, cells with 2, 3 or 4 SPBs appeared slightly less than 1 h before cells with the same number of nuclei, with 96% of cells completing two SPB divisions by 9 h post induction of meiosis (Fig. 2b, c). In the *srs2-101* strain, cells that had accomplished SPB division and separation increased with wild-type timing until 6 h and then increased only slightly. By 9 h, more than 75% of cells had produced at least 2 separated SPBs, while only 60% of cells had completed the meiosis I nuclear division (Fig. 2b), and about 35% of cells had failed to complete the two SPB and nuclear divisions (Fig. 2c). Most notably, a significant population of *srs2* cells had divided SPBs but contained only an undivided nuclear signal. The fraction of cells with this phenotype reached a maximum at 6 h, with 32% and 24% of *srs2-101* and *srs2Δ* cells, respectively, compared to 8.0% in wild-type (Fig. 2e). This observation, that loss of Srs2 activity results in a significant population of cells with divided SPBs but undivided nuclei, suggests that these cells are attempting to progress through meiosis II despite the nucleus failing to divide correctly in meiosis I. However, it should also be noted that about a quarter of the population does not undergo even the first round of meiotic SPB separation.

**Fig. 2.**
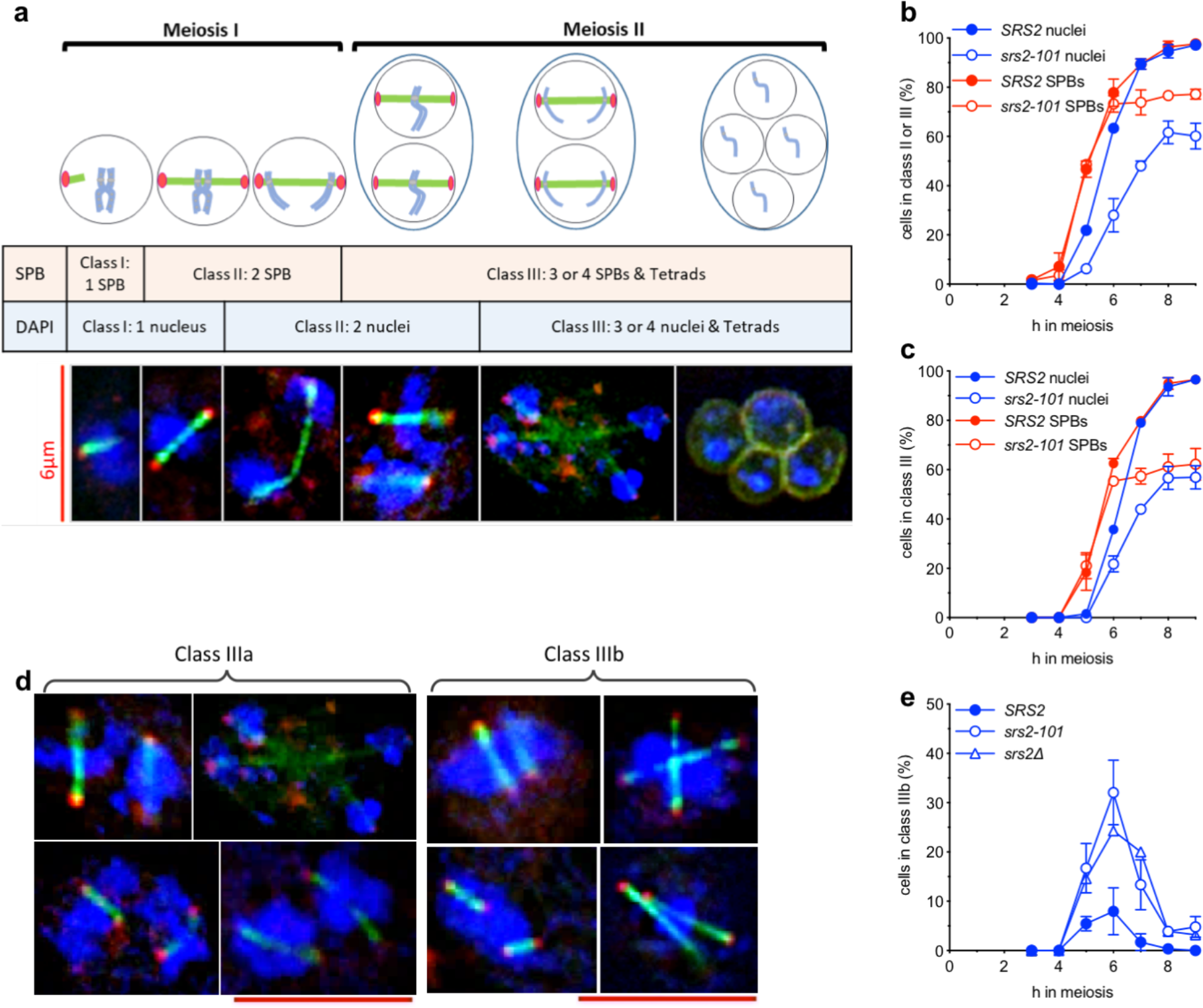
Altered nuclear division and spindle dynamics in the absence of Srs2 activity. (b) Classifications used during analysis of spindle pole body (SPB) and nuclear (DAPI) division. (c) Fraction of cells completing both meiotic nuclear (blue) or SPB divisions. Unlike nuclear division (see also Fig. 3), SPB division is not delayed in *srs2-101* [nuclear division as in panel (a); SPB analysis, n=2; error bars—range]. (e) Combining nuclear and SPB analysis reveals two distinct populations of cells that have completed both SPB divisions: Class IIIa, cells with at least two nuclei, as expected for cells with divided SPBs; and Class IIIb, cells with only a single nucleus, despite having divided SPB signals. Scale bar = 6μm. (f) Loss of Srs2 activity is associated with an increased frequency of class IIIb nuclei, which have completed SPB division but have failed to divide nuclei (*SRS2*, n=2; *srs2-101*, n=2; *srs2*Δ, n=1. error bars—range)

### Sister centromeres separate in the absence of Srs2, despite failures in nuclear division

To investigate the nature of the nuclear division failure in *srs2* strains, we monitored sister chromatid separation, using strains that expressed a fusion between the Tet repressor and GFP (TetR-GFP) and that were heterozygous for a Tet operator array (*tetO*, 224 repeats) inserted near the centromere of chromosome V. In such strains the separation of the *tetO*/TetR signals is a proxy for sister centromere separation. In wild-type cells, a single signal is visible until the meiosis II, when this signal divides into two signals, which then segregate into two of the four nuclei present at the tetranucleate stage (Fig. 3a).

**Fig. 3.**
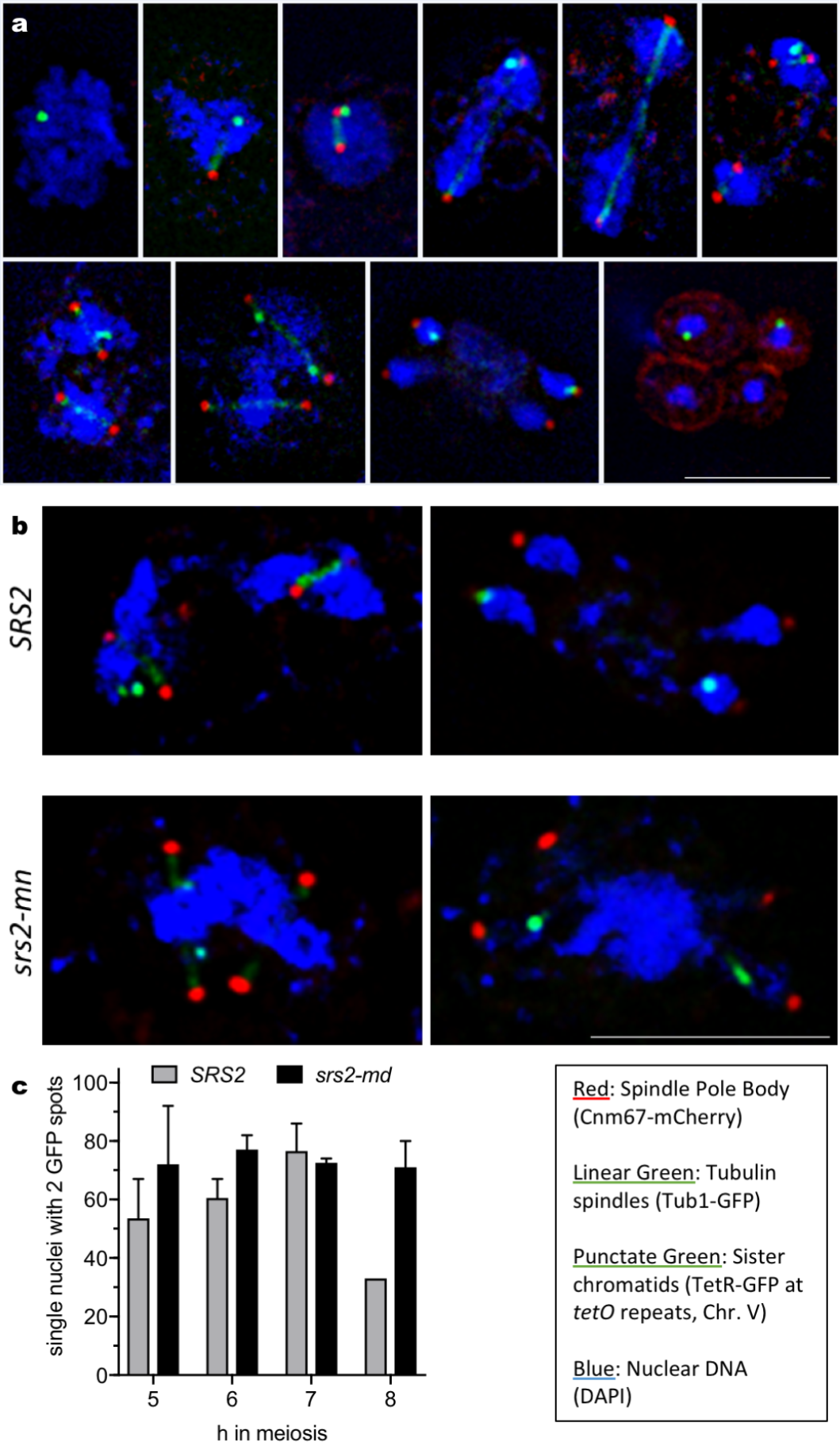
Sister chromatid separation observed during meiosis using a tet repressor-GFP fusion and heterozygous tet operon repeats at the *URA3* locus. (a) Representative images showing progression of sister chromatid segregation through meiosis in wild-type cells. Scale bar = 5μm. (b) Representative images of *SRS2* (6h & 7h) and *srs2-md* (6h & 8h) cells with divided spindle pole bodies. Even when the nucleus fails to divide, separated *tetO*/TetR signals can be observed suggesting sister chromatids have separated. Scale bar = 5μm. (c) Cells with divided signals as a percentage of those cells with a single nucleus but divided SPBs (i.e. Class IIIb cells, see Fig. 4e). n=2, except for *SRS2* at 8h for which Class IIIb cells were only observed in one experiment; error bars denote range.

Cells with divided SPBs but one nucleus (i.e. Class IIIb cells, see Fig. 2d) were analysed for *tetO*/TetR signal separation. Interestingly, most Class IIIb cells in the *srs2-md* strain had two *tetO*/TetR signals, with percentages similar to wild-type, despite the nucleus failing to divide (Fig. 3b, c). Although fewer wild-type cells displayed single nuclei at late time points in meiosis, it is notable that the *tetO*/TetR signals in these cells also usually separated. This suggests that even when nuclear division fails, the majority of sister centromeres are separating correctly, at least where measured close to pericentromeric DNA on Chromosome V. Similar analysis of a strain with homozygous *tetO* inserts also indicates that 4 SPB signals are frequently observed in *srs2* strains (EH and ASHG, unpublished observations), suggesting that centromeres on both homologous chromosomes and sister chromatids separate.

### Loss of Srs2 function in meiosis leads to Rad51 aggregate formation

As Srs2 is thought to regulate mitotic recombination by removing Rad51 from ssDNA nucleoprotein filaments and allowing repair by SDSA (Andersen and Sekelsky, 2010), we hypothesized that the meiotic nuclear division failure seen in *srs2* cells may be due to entanglements caused by a failure to remove Rad51. We therefore analysed Rad51 distributions in nuclear spreads using immunofluorescence. Cells were classed into three groups: those with no Rad51 signal; those with only small Rad51 signals (<0.39μm in any direction), hereafter called ‘Rad51 foci’; and those with large Rad51 signals (>0.39 μm), which we here refer to as ‘Rad51 aggregates’ (Fig. 4a). *srs2* mutants displayed persistent Rad51 signal (Fig. 4b) most of which was due to a significant increase in the proportion of cells with Rad51 aggregates, which were rarely seen in wild-type cells (Fig. 4d). A substantial fraction of *srs2* cells also displayed persistent Rad51 foci, at times when they had disappeared from wild type (Fig 4c).

**Fig. 4.**
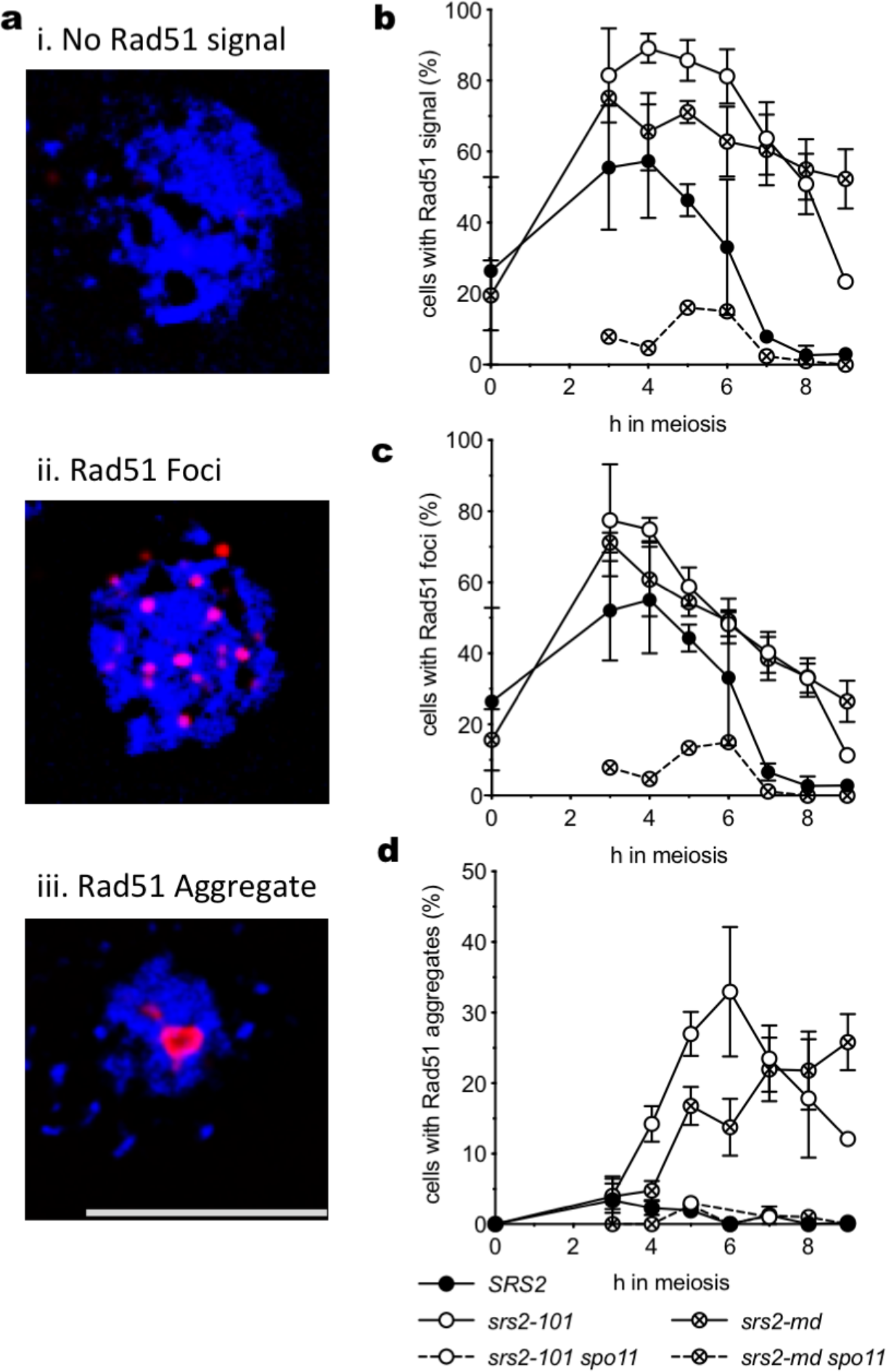
Mutants lacking Srs2 activity form Rad51 aggregates. (a) detection of Rad51 in nuclear spreads; blue=DAPI/DNA, red=α-Rad51. Cells were categorised as: i. cells with no Rad51 signal; ii. cells with Rad51 foci, iii. cells with large Rad51 aggregates. Scale bar = 5μm. (b) *srs2* mutants show a persistent Rad51 signal (either foci or aggregates). (c) *srs2* mutants display Spo11-dependent Rad51 foci. (d) *srs2* mutants display Spo11-dependent Rad51 aggregates. (*SRS2*, n=3; *srs2-101*, n=3; *srs2-md*, n=5; *srs2-101 spo11*, n=1; *srs2-md spo11*, n=1; error bars—standard error of the mean).

Meiotic DSB formation requires Spo11 transesterase activity, which is abolished by loss of the reactive OH group at the catalytic tyrosine (*spo11-Y135F*). Both aggregate formation (Fig. 4b, c) and sporulation defects (Fig. S2a) of *srs2* mutants were suppressed by the *spo11-Y135F* mutation, suggesting that both Rad51 aggregates and barriers to nuclear division form as a consequence of abnormal DSB repair in the absence of Srs2.

### Meiotic phenotypes of srs2 depend on Rad51–mediated strand invasion

Rad51 protein has two activities with the potential to contribute to aggregate formation during meiosis in *srs2* mutants. Rad51 forms a filament on ssDNA, which then catalyzes invasion of homologous duplex DNA through a second DNA binding site. *rad51-II3A* mutants, which lack this second binding site, are competent for filament formation but defective in strand invasion (Cloud et al., 2012). During meiotic recombination in budding yeast, Rad51 acts as a co-factor in filament assembly but not strand invasion with a meiosis-specific homologue, Dmc1, and *rad51-II3A* mutants are competent for meiotic DSB formation and homologous recombination (Cloud et al., 2012). If the phenotypes observed in *srs2* mutants were due to a defect in removing Rad51 from normal meiotic recombination intermediates, then aggregates should still form in *srs2 rad51-II3A* double mutants. Because *rad51-II3A* and *srs2-101* or *srs2Δ* mutants are synthetically lethal (T-CC, AH and ASHG, unpublished observations), we combined the meiotic-depletion *srs2-md* allele with *rad51-II3A*. Only ~4% of *srs2-md rad51-II3A* cells contained Rad51 aggregates, as compared to 15-25% of *srs2-md RAD51* cells (Fig 5b). This indicates that aggregate formation depends on Rad51’s ability to bind a second DNA molecule, rather than retention of Rad51 filiaments. Since Rad51 strand exchange activity is not normally required for meiosis, this suggests that Rad51 aggregates derive from “off-pathway” events, rather than from normal meiotic recombination intermediates.

**Fig. 5.**
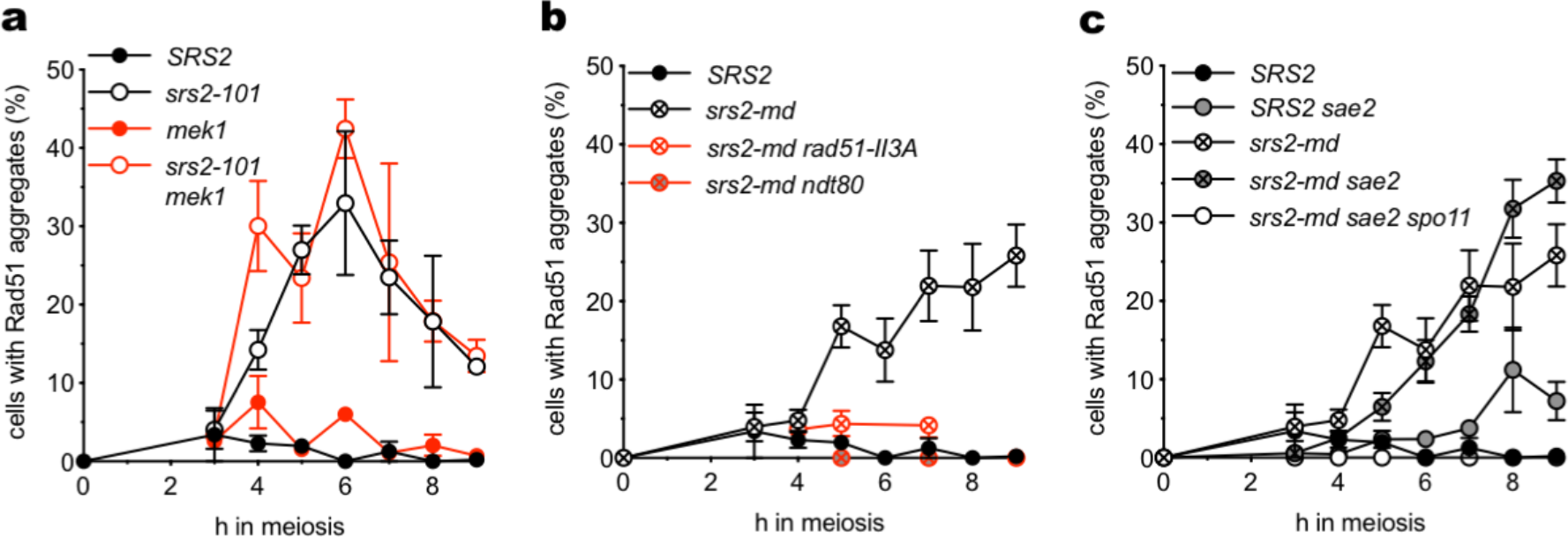
Analysis of the Rad51 aggregation phenotype. (a) Deletion of *MEK1* does not suppress aggregate formation, indicating that an inter-homologue bias for strand invasion is not required. (b) Rad51 aggregates do not form in *rad51-II3A* or *ndt80*Δ mutants, indicating that both Rad51-mediated strand invasion and exit from pachytene are required for aggregate formation. (c) Deletion of *SAE2* does not prevent aggregate formation, suggesting that normal DSB processing is not required. Interestingly, a small fraction of *sae2Δ* cells form aggregates at later time points in a *SPO11*-dependent manner. (*SRS2*, n=3; *srs2-101*, n=3; *srs2-101 mek1Δ*, n=2; *mek1Δ*, n=2; *srs2-md*, n=5; *srs2-md ndt80Δ*, n=1; *srs2-md rad51-II3A*, n=3; *srs2-md sae2Δ*, n=4; *sae2Δ*, n=3; *sae2Δ spo11-Y135F*, n=1). Error bars—standard error of the mean, except for *srs2-101 mek1*Δ and *mek1*Δ, where they denote range.

### Rad51 aggregation is independent of MEK1-dependent interhomologue bias

To test whether Rad51 aggregates that form in *srs2* mutants arise from normal interhomologue meiotic recombination intermediates, we examined mutants lacking Mek1, a meiosis-specific kinase that is a major effector of the bias towards use of the homologue, rather than the sister chromatid, during meiotic DSB repair (Niu et al., 2005). Deletion of *MEK1* did not rescue the aggregation phenotype of *srs2-101* mutants, suggesting Rad51 aggregates are not caused by a failure in the processing or resolution specifically of interhomologue recombination intermediates (Fig. 5a).

### Rad51 aggregate formation is independent of Sae2

As Rad51 aggregate formation depends on Spo11-DSB formation and the ability of Rad51 to bind a second DNA molecule, we hypothesized that Rad51 aggregation should also be dependent on DSB resection. To test this, we performed analysis in cells lacking Sae2, which is required for normal resection of Spo11-induced DSBs prior to strand invasion. Surprisingly, however, deletion of *SAE2* did not prevent the formation of aggregates, which at late time points were instead slightly increased in *srs2-md sae2Δ* double mutants compared to *srs2-md* alone (Fig. 5c). Rad51 aggregates were also observed in *sae2Δ SRS2* cells at later time points, and these aggregates were also Spo11-dependent. The proportion of aggregates seen at late time points in *srs2-md sae2Δ* double mutants reflected the sum of levels seen in the single mutant cells, consistent with an independent contribution from the two single mutations (Fig. 5, c). Thus, Rad51 aggregate formation does not require normal meiotic DSB resection.

### Rad51 aggregates form after exit from pachytene

Since analysis of *srs2 rad51-II3A* and *srs2 mek1* double mutants suggested that Rad51 aggregates form as the consequence of abnormal meiotic recombination, it was of interest to determine the stage of meiosis at which aggregation occurs. Rad51 immunofluorescence was combined with visualisation of the synaptonemal complex (SC) using GFP-tagged Zip1 (White et al., 2004), a component of the tripartite structure that forms along the length of homologous chromosomes during the pachytene stage of meiotic prophase and provides a framework that assists the process of homologous recombination (Heyting, 1996; Yang and Wang, 2009). Once recombination is completed, cells exit pachytene in a process that requires gene expression driven by the Ndt80 transcription factor (Chu and Herskowitz, 1998). The SC is disassembled and recombination intermediates resolve, leaving the crossover products of recombination to maintain connections between homologues (Sourirajan and Lichten, 2008; Zickler and Kleckner, 1999, 2016). As expected, Rad51 foci were found in wild-type cells with either Zip1 foci or long regions of synapsis, but were absent from cells lacking Zip1 signal. However, *srs2* mutants displayed both Rad51 foci in Zip1-positive cells, and prominent Rad51 aggregates in cells that lacked any Zip1 signal, suggesting that aggregates form after cells exit pachytene (Fig. 6). To confirm this, we examined *ndt80Δ* mutants, which arrest in pachytene with fully synapsed homologues and duplicated but unseparated SPBs (Xu et al., 1995). Rad51 aggregates were absent from *srs2-md ndt80Δ* cells at all times (Fig. 5b). These results indicate that, in *srs2* mutants, Rad51 aggregate formation occurs after the cells exit from pachytene, and thus after meiotic recombination would be complete in wild-type cells.

**Fig. 6.**
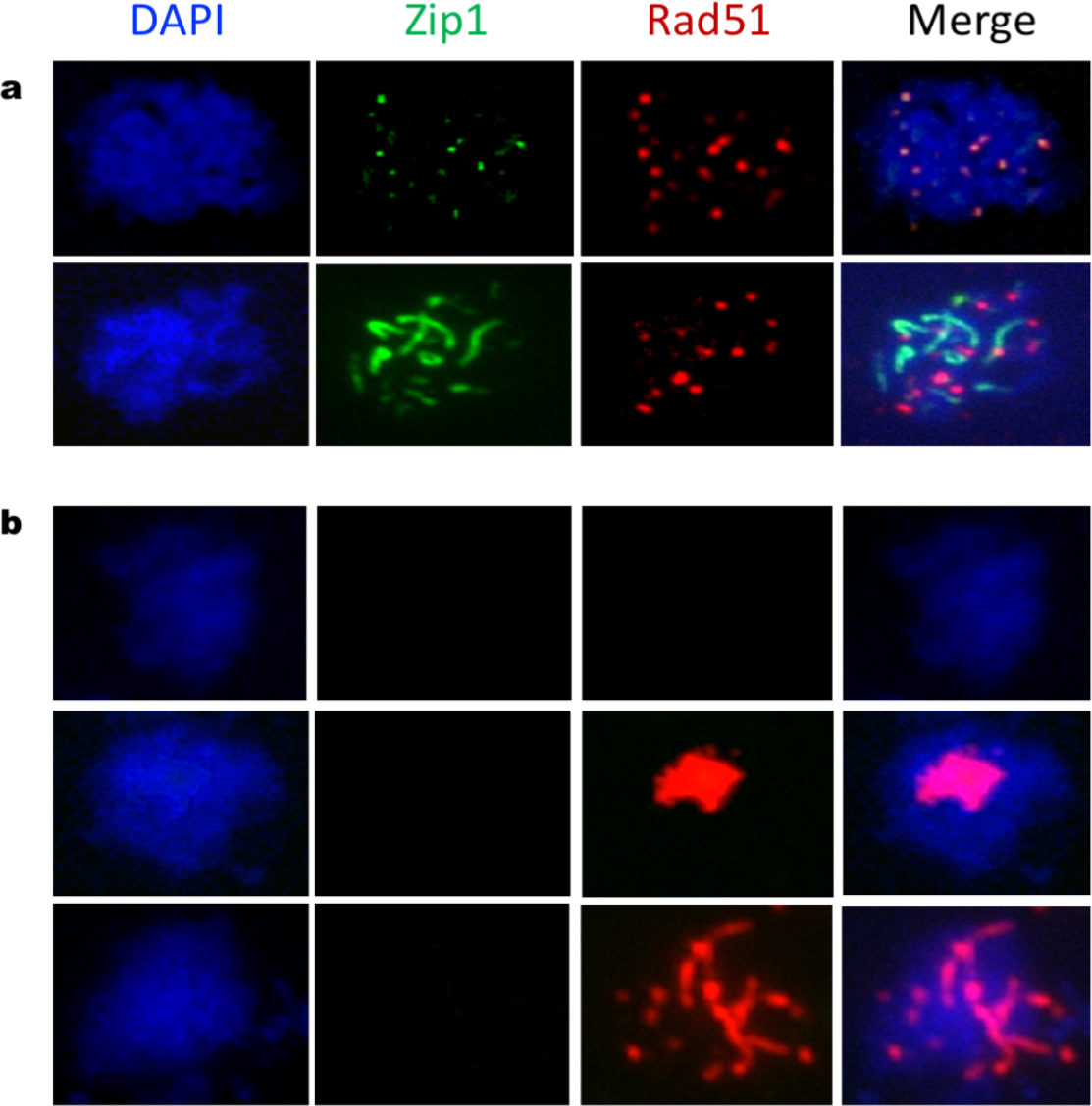
Analysis of Rad51 aggregates in cells expressing Zip1-GFP, a synaptonemal complex protein. (a) Representative images of *srs2-101* cells with punctate or elongated Zip1-GFP signals (3 h and 5 h, respectively) with clear Rad51 foci. (b) Representative images of Zip1-negative *srs2-101* cells at 5 h. Cells can be classified as having no Rad51 signal, large aggregates of Rad51 or elongated thread-like aggregates of Rad51. Aggregates of Rad51 can be observed in *srs2-101* cells in the absence of a synaptonemal complex.

### RPA colocalizes with Rad51 aggregates

The ssDNA that is formed by DSB resection is initially coated with Replication Protein A (RPA), a heterotrimeric complex that is essential for viability (Brill and Stillman, 1991; Chen et al., 2013). Although RPA binding to ssDNA is important for formation of active Rad51 filaments, RPA is displaced during filament formation. Thus, Rad51 and RPA foci are at times adjacent to each other, but rarely are truly colocalized (Gasior et al., 1998; Hays et al., 1998). However, a substantial fraction of Rad51 aggregates that form in *srs2-md* colocalize with either a focus or an aggregate of RPA (Fig. 7a, b). Detailed analysis of Rad51 and RPA signals at 5 h post induction of meiosis in *srs2-md* cells confirmed that foci of RPA and Rad51 generally do not colocalize, but Rad51 and RPA are frequently colocalized in aggregates, with more than 90% of Rad51 aggregates colocalizing with RPA (Fig. 7c), consistent with the suggestion that the Rad51 aggregates that form in *srs2* mutants reflect the presence of abnormal recombination events.

**Fig. 7.**
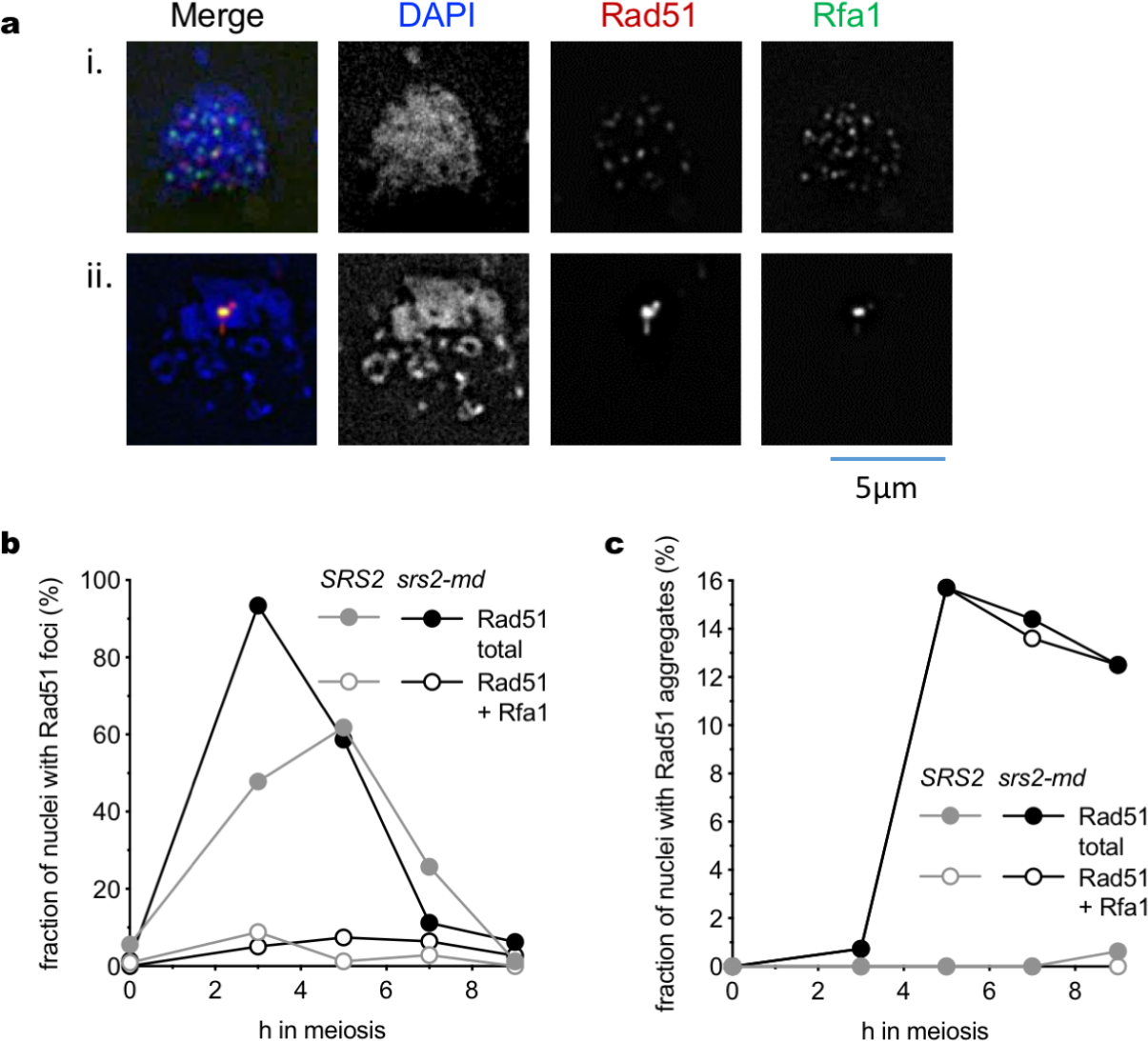
Rad51 aggregates frequently colocalize with RPA. (a) Representative images of: i. a nucleus with both Rad51 and Rfa1 foci, with minimal colocalization; ii. a Rad51 aggregate colocalized with Rfa1. (b) RPA colocalization with Rad51 foci across a time course. Colocalization between Rad51 foci and Rfa1 is infrequent. c) RPA colocalization with Rad51 aggregates in the same experiment. Almost all Rad51 aggregates that form in *srs2-md* cells colocalize with either foci or aggregates of Rfa1. >100 nuclei were scored for all time points, except for *SRS2*, 7h and *srs2-md*, 0h (70 and 72 nuclei, respectively).

### Altered recombination intermediates in srs2 mutants

The above analysis indicates that aggregates of Rad51 do not form until cells exit from pachytene, which is also the time that double Holliday junction (dHJ)-containing recombination intermediates (joint molecules; JMs) are resolved as COs (Allers and Lichten, 2001; Sourirajan and Lichten, 2008). To test the suggestion that Rad51 aggregate formation might be associated with abnormal JM metabolism, molecular analysis of recombination intermediates and products was performed, using a DNA isolation method that preserves dHJ intermediates (Allers and Lichten, 2000). Southern blots were used to detect events at a *URA3-ARG4* recombination reporter (Jessop et al., 2005) inserted on the left arm of chromosome III (Jessop et al. 2005)(Fig. 8, Supplementary Fig. S3a). While DSB levels were similar in *srs2* and *SRS2* strains (Supplementary Fig. S3b), a marked deficit in levels of detectable JMs (4 to 7-fold) was observed in *srs2* mutants (Figure 8a, b). Despite the apparent reduction in JMs, which are crossover precursors (Allers and Lichten, 2001), crossovers were recovered at similar levels from wild-type and *srs2* mutant cells (Figure 8c).

**Fig. 8.**
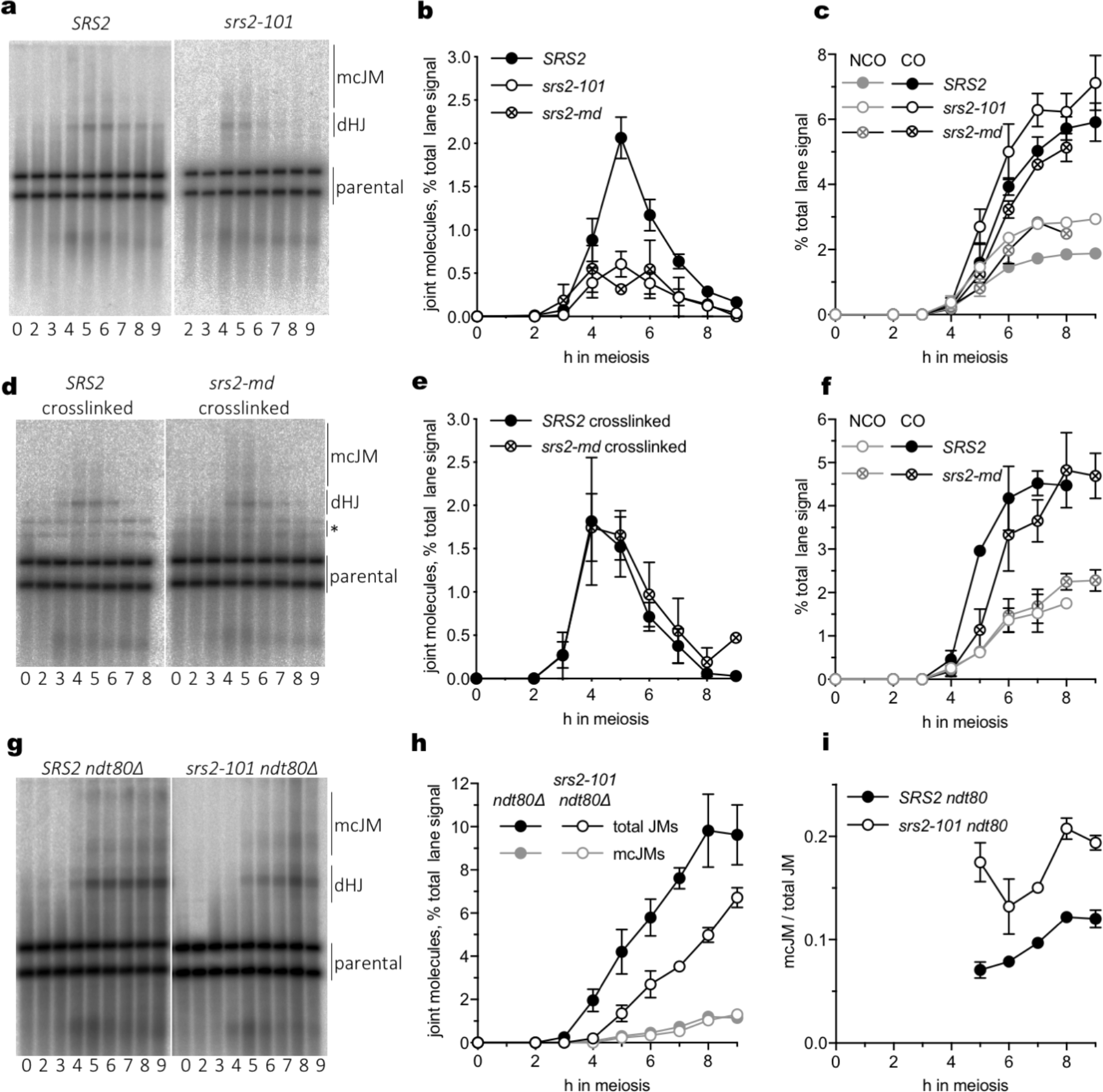
Differences between recombination intermediates (JMs) formed in the presence and absence of Srs2 activity. (a) Southern blots of *Xmn*I digests of DNA from the indicated time after initiation of meiosis, probed to detect joint molecules (dHJ—double Holliday junctions; mcJM, multichromatid joint molecules) at a recombination reporter on chromosome *III* (see Figure S1). (b) quantification of blots from *SRS2* (n=5, data from (De Muyt et al., 2012; Kaur et al., 2015) and one additional replicate; error bars denote S.E.M.), *srs2-101* (n=5; error bars denote S.E.M.) and *srs2-md* (n=2; error bars denote range). JM recovery is markedly reduced in the absence of Srs2 activity. (c) Crossovers are formed at similar levels in the presence and absence of Srs2 activity, despite the marked difference in JM recovery. Noncrossovers are recovered at ~50% greater levels in *srs2* mutants than in wild type. (d) Detection of JMs, as in panel (a), in psorlaen-crosslinked DNA samples. Bands marked with an asterisk are the products of crosslink-induced partial *XmnI* digestion. (e) quantification of 3 gels from two sporulations; error bars denote S.E.M. Psoralen crosslinking restores parity in JM recovery to *srs2* mutants. (f) Crossovers and noncrossovers from crosslinked samples. (g) Detection of JMs, as in (a), from JM resolution-defective *ndt80* mutants. (h) quantification of gels. n=2; error bars denote range. Total JM appearance in *srs2* mutants is delayed and reduced ~ 2-fold, but mcJMs are similar in *SRS2* and *srs2*. (i) fraction of JMs that are mcJMs.

We considered two possible explanations for how steady state levels of JMs could be reduced so markedly without substantially affecting CO levels. First, JM resolution might be accelerated, such that the average JM lifespan in *srs2* mutants was 4 to 8 times shorter than in wild type. Second, JMs formed in *srs2* mutants might contain structural differences (such as nicked Holliday junctions) that make them unstable, either *in vivo* or during the DNA purification protocol used here. To test the first suggestion, JMs were measured in DNA isolated from resolution-defective *ndt80Δ* mutants using the same protocol, and were found to be both delayed and substantially reduced in *srs2 ndt80Δ* relative to *SRS2 ndt80Δ*. Thus, at least some of the difference between *SRS2* and *srs2* must be due to factors that affect JM structure or stability, rather than resolution. Consistent with this suggestion, loss of Srs2 activity resulted in a substantial increase (1.6-fold, average of 6-9h) in the fraction of JMs in resolution-defective *ndt80Δ* mutants that contained more than two chromatids (multichromatid JMs; Figure 8).

To test the second suggestion of increased lability of JMs formed in *srs2* mutants, the DNA isolation protocol was modified to include psoralen crosslinking, which has the potential to preserve otherwise unstable intermediates (Kaur et al., 2018). The addition of psoralen crosslinking resulted in recovery of JMs in similar levels from *SRS2* and *srs2* strains (Figure 10e), supporting the suggestion that Srs2 activity is important for normal JM formation and stability.

## Discussion

While multiple studies have examined roles for the multifunctional *S. cerevisiae* helicase Srs2 in DNA replication, repair and recombination during the mitotic cell cycle (Niu and Klein, 2017), considerably less is known about Srs2’s function during meiosis. Previous studies had shown that loss of Srs2 activity causes delays and defects in meiotic nuclear division, as well as reduced sporulation and spore viability (Palladino and Klein, 1992; Sasanuma et al., 2013b). Our current study and an accompanying study (Sasanuma et al., 2019) have identified recombination abnormalities in meiosis I prophase that are likely the cause of abnormal nuclear division in *srs2* mutants.

We find that *srs2* mutants display multiple meiotic defects at both the cellular and molecular level, regardless of whether the mutants examined confer a chronic loss of Srs2 activity (*srs2*Δ and *srs2-101*) or a loss of Srs2 specifically during meiosis (*srs2-md*). These and other *srs2* mutant phenotypes are Spo11-dependent [Fig. S2a, Fig. 4, Sasanuma et al. (2019)], indicating that abnormal recombination intermediates are a likely cause. A minority of *srs2* mutant cells (21-27%) do not divide their nuclei (Fig. 1) and do not separate SPBs (Fig. 2c), consistent with a DNA damage-induced meiotic progression arrest before exit from pachytene. Cells that do exit from pachytene and progress to nuclear division do so with an ~1 h delay (Fig. 1), even though SPB separation occurs with normal timing during meiosis I and meiosis II (Fig. 2b, c). These cells display DNA bridges (Fig. 1f), sister centromere separation without nuclear division (Fig. 3), meiosis II SPB separation and spindle formation within a single nuclear mass (Fig. 2d, e), and substantial spore inviability (Fig. 1g, h). These latter phenotypes suggest frequent recombination-dependent chromosome entanglements preventing nuclear division, as is observed in other mutants that are unable to properly resolve recombination intermediates (De Muyt et al., 2012; Jessop and Lichten, 2008; Kaur et al., 2015; Oh et al., 2008; Tang et al., 2015). In summary, the pleotropic nature of the meiotic nuclear division defects seen in *srs2* mutants are consistent with abnormal recombination intermediates being present both before and after cells exit from pachytene.

### Persistent abnormal recombination intermediates in srs2 mutants

Srs2 is thought to disrupt or prevent formation of Rad51-containing nucleoprotein filaments during the mitotic cell cycle (Niu and Klein, 2017). The observation of persistent Rad51 aggregates in *srs2* mutants (Figure 4; Sasanuma et al., 2019) suggests that Srs2 has a similar function during meiosis. Interestingly, while aggregate formation does not require progression through meiosis I (Sasanuma et al., 2019), the absence of aggregates from *ndt80*Δ mutants, and from *NDT80* cells with foci or linear arrays of the SC protein Zip1 (Figs. 5b and 6; Sasanuma et al., 2019), indicates that Rad51 aggregates do not form until cells have exited from pachytene and disassembled SC. Rad51 strand exchange activity is inhibited before pachytene exit (Subramanian et al., 2016), and Rad51 aggregate formation is greatly reduced in *srs2 rad51-II3A* mutants (Fig. 5b), which can form Rad51-DNA filaments but not catalyse strand exchange activity (Cloud et al., 2012). Thus, the appearance of Rad51 aggregates after exit from pachytene indicates that Rad51 strand exchange activity is necessary for this *srs2* mutant phenotype. Notably, the frequent colocalization of aggregates of Rad51 and the RPA subunit Rfa1, observed frequently in *srs2* mutants but rarely during normal meiosis (Fig 7), suggests these aggregates form on abnormal recombination intermediates, most likely associated with stalled recombination events that are impeding normal nuclear division.

We applied two tests to determine if the Rad51 aggregates were likely to be associated with normal or abnormal recombination events. First, we deleted the Mek1 kinase, which during normal meiosis directs repair of Spo11-DSBs away from the sister chromatid and toward the homologue (Callender et al., 2016; Niu et al., 2005; Niu et al., 2009). Secondly, we deleted Sae2, which is required for resection of Spo11-DSB ends during meiosis I prophase (McKee and Kleckner, 1997; Prinz et al., 1997). Rad51 aggregation still occurred in *mek1*Δ *srs2* and in *sae2*Δ *srs2* (Fig. 5a, c), indicating that neither interhomologue strand invasion nor Sae2-mediated single-strand resection are required for post-pachytene Rad51 aggregation in *srs2* cells. This adds further support to our hypothesis that the cytologically visible phenotypes of *srs2* mutants might reflect the presence of abnormal recombination intermediates after exit from pachytene. It leaves open the question of whether these abnormal intermediates arise as a consequence of *srs2* mutant defects during early meiosis I prophase, when recombination normally occurs, or as a consequence of aberrant processing of residual lesions present during late meiosis I prophase, after exit from pachytene and the time when most DSB repair is complete.

### Early defects in recombination intermediate formation in srs2 mutants

Analysis of recombination intermediates at a *URA3-ARG4* recombination reporter revealed that, DSBs are formed and repaired with normal kinetics in *srs2* mutants (Fig. S3b), and both crossover and noncrossover recombinants were formed in *srs2* mutants at levels approximating those seen in wild-type. Thus, in contrast to what is seen in studies of vegetatively growing cells, Srs2 does not appear to have either pro- or anti-crossover function during meiosis. However, JMs, which are crossover precursors, were recovered from *srs2* mutants at reduced levels when DNA was extracted using a protocol that normally stabilizes JMs, but at normal levels when DNA was psoralen-crosslinked before extraction (Fig. 8). Loss of Srs2 activity results in both delays and reductions in recovery of stable JMs accumulating in resolution-defective *ndt80* cells, and the proportion of multichromatid JMs, which form when DSB ends interact with more than one repair template (Oh et al., 2007), is also increased. These findings suggest that many of the JMs formed in *srs2* mutants have less stable structures than JMs formed in wild type cells, possibly reflecting a failure to form fully ligated double Holliday junctions. These unstable JMs may be derived from a yet uncharacterized Srs2-dependent, Dmc1-independent pathway for meiotic DSB repair, suggested by observations that more unrepaired DSBs accumulate in *srs2 dmc1* double mutants than in *SRS2 dmc1* (T-CC and ASHG, unpublished observations).

### Concluding remarks

Taken together, our data and the data of Sasanuma et al. (Sasanuma et al., 2019) indicate that Srs2 has an important role in recombination biochemistry throughout meiosis I prophase. We propose that a major function of Srs2 during meiosis is to limit residency of Rad51 on Spo11-initiated DSBs. In the absence of Srs2 activity, increased Rad51 occupancy may reduce the ability of DSB ends to complete dHJ formation, both because fewer binding sites are available for Dmc1 filament formation, and because Rad51 strand invasion activity is inhibited during early meiosis I prophase (Busygina et al., 2008; Callender et al., 2016; Niu et al., 2009; Tsubouchi and Roeder, 2006). When *srs2* mutant cells exit from pachytene, relief of this inhibition (Subramanian et al., 2016) might allow transient formation of fully ligated dHJ intermediates before resolution as crossovers; alternatively, recombination intermediates formed in *srs2* mutants may not be fully ligated, but can still can be resolved as COs upon exit from pachytene (Whitby, 2005). We also suggest that, in *srs2* mutants, Rad51 aggregates that form upon exit from pachytene reflect the persistence of lesions that seed further Rad51 accumulation in the absence of Srs2 translocase; whether these are aberrant events normally resolved by Srs2, unresolvable intermediates formed before exit from pachytene, or intermediates formed *de novo* by DSBs present after pachytene exit, remains to be determined. The frequent, Spo11-dependent formation of Rad51 aggregates in *srs2 sae2* mutant cells (Fig. 5c), which are not expected to undergo resection or strand invasion before exit from pachytene, most likely reflects such post-pachytene DSB processing, either through DSB end-destabilization by high Rad51 residency, or through end-resection by nucleases or other activities that are derepressed after pachytene exit.

Srs2 belongs to the highly-conserved UvrD helicase family, which has at least one representative in most organisms (Lorenz, 2017; Niu and Klein, 2017). Loss of the Srs2-like Fbh1 protein in *Schizosaccharomyces pombe*, which also has Rad51 removal activity, confers meiotic defects similar to those seen in budding yeast, including reduced spore viability, meiotic nuclear division defects, and persistent Rad51 foci, without affecting frequencies of crossovers or noncrossovers among viable progeny (Sun et al., 2011). Interestingly, the human Fbh1 orthologue, when expressed in budding yeast, suppresses many of the mitotic recombination defects and DNA damage sensitivity of *srs2* mutants, suggesting that hFBH1 may have Rad51-removing activity, as well (Chiolo et al., 2007). It will be of considerable interest to determine the molecular defects underlying the similar mutant phenotypes of budding yeast *srs2* and fission yeast *fbh1*-mutants, and to determine if the verterbrate protein has similar functions.

## Materials and Methods

### Yeast strains

All yeast strains (Supplementary Table S1) are derived from SK1 (Kane and Roth, 1974) by either transformation or genetic crosses. The *srs2-md* meiotic depletion allele (*pCLB2-3HA-SRS2*) inserts CLB2 promoter sequences and a triple influenza hemagglutinin epitope tag immediately upstream of *SRS2* coding sequences, and was constructed by amplification of *pCLB2-3HA* sequence from pAG335 (a gift from Monica Boselli) with primers containing 50 nt targeting the *SRS2* locus: Srs2Clb2ptagF—GAGTATCATTCCAATTTGATCTTTCTTCTACCGGTACTTAGGGATAGCAAtcgatgaattcgagctcg; Srs2CClb2ptagR: GTATT TAACTGGGATACTAAATGCAACCAAAGATCATTGTTCGACGACATgcactgagcagcgtaatctg, where lowercase letters correspond to plasmid sequences. *CNM67-mCherry* and *GFP-TUB1* tag constructs were from AMY6969 &AMY6706, respectively (a gift from Adele Marston). Zip1-GFP has been described (White et al., 2004). The *RFA1-GFP* allele inserts *GFP* sequences immediately downstream of *RFA1* coding sequences, and was constructed by amplification of *GFP* sequence from pKT127 (Sheff and Thorn, 2004) with primers containing 50 nt targeting the RFA1 locus: RFAChrGFP_F: GGGCTGAAGCCGACTATCTTGCCGATGAGTTATCCAAGGCTTTGTTAGCTggtgacggtgctggttta;RFAChrGFP_R:TTTCTCATATGTTACATAGATTAAATAGTACTTGATTATTTGATACATTAtcgatgaattcgagctcg, where lowercase letters correspond to plasmid sequences.

### Spore viability

Spore viability was determined by tetrad dissection. Diploid strains were sporulated 1-3 days, 30°C in patches on 1% (w/v) potassium acetate, 2% (w/v) agar plates, digested 15 min, 30°C in 1M sorbitol, 1% (w/v) glucose, 1 mg/ml zymolyase, dissected on YPAD agar (2% peptone, 1% yeast extract, 2% glucose, 0.004% adenine, 2% agar, all w/v) and incubated for 2 days at 30°C.

### Liquid sporulation

All incubation temperatures were 30°C. All media components are w/v unless otherwise indicated. For cytological studies, an overnight YPAD broth culture, inoculated with a single colony on YPAD agar, was grown with aeration overnight, and then was diluted to an OD_600_ of ~0.3 in BYTA (1% yeast extract, 2% tryptone, 1% potassium acetate, 50 mM potassium phthalate) and incubated with aeration for 16 h. The culture was harvested by centrifugation, washed with 200 ml 1% potassium acetate, resuspended in 250 ml SPM (0.3% potassium acetate, 0.02% raffinose) with appropriate supplements to a final OD_600_ of ~1.9, and incubated with vigorous aeration. For molecular studies, sporulation in liquid was as described (Oh et al., 2009), using the “Lichten lab” protocol.

### Cytological methods

#### Nuclear divisions

0.5 ml of a culture was mixed with 0.75 ml ethanol and stored at −20°C. 1μl 0.5 mg/ml 4’,6-diamidino-2-phenylindole (DAPI) was added and samples were incubated at room temperature for 30 sec. Cells were harvested by centrifugation, resuspended in 0.2 ml 50% glycerol, sonicated briefly if necessary to break up clumps, and examined by epiflouresence microscopy. Cells with 2 nuclei were scored as having completed meiosis I; cells with 3 or 4 nuclei as having completed meiosis II.

#### Recombination protein foci

4.5 ml of a culture was harvested by centrifugation and resuspended to 0.5 ml in 1.0 M sorbitol, pH 7. 1,4-dithiothreitol and zymolyase were added to 24 mM and 0.14 mg/ml, respectively, and cells were spheroplasted by incubation at 37°C for 20 to 45 min with agitation, until the majority of cells lysed when mixed with an equal volume of 1.0% (w/v) sodium N-lauroylsarcosine. 3.5 ml of Stop Solution (0.1 M MES, 1 mM EDTA, 0.5 mM MgCl_2_, 1 M sorbitol, pH 6.4) was added, cells were harvested by centrifugation, resuspended in 100 μl Stop Solution, and distributed between four ethanol-cleaned glass microscope slides. 20 μl of fixative [4.0% (w/v) formaldehyde, 3.8% (w/v) sucrose, pH7.5] was added to each slide, followed by 40 μl of 1% Lipsol with light mixing. A further 40 μl of fixative was added, and the mixture was spread across the slide. All subsequent steps were at room temperature, unless otherwise indicated. Slides were incubated for 30 min in a damp chamber, and then allowed to air dry. Once dry, slides were washed in 0.2% (v/v) PhotoFlo (Kodak) and then in water and allowed to air-dry slightly. Slides were washed once in 0.025% Triton X-100 in phosphate-buffered saline (PBS) for 10 min and twice for 5 min in PBS. Slides were blocked with 5% skimmed milk (Sigma) in PBS for 1-4 h at 37°C, excess liquid was removed and slides were placed horizontally in a damp chamber. Mouse anti-yeast Rad51 antibody (Santa Cruz, sc-133089, 1:200 in 1% skim milk, PBS) was added at 150 μl per slide and slides were incubated at 4°C overnight. Slides were washed three times in PBS (5 min each) and incubated with secondary antisera (AlexaFluor594-conjugated goat anti-mouse antibody;Life Technologies A11005; 1:1000 in 1% skimmed milk, PBS), 150 μl per slide, for 1-2 h in a damp chamber. Slides were washed three times with PBS (5 min each), cover slips were affixed using Vectashield mounting medium with DAPI (Vector Laboratories), sealed with clear varnish and imaged on a DeltaVision microscope (z=12-15, Exposure times: RD-TP-RE=1.0 s, DAPI=0.05-1.0 s, FITC = 1.0 s for Rfa1-GFP colocalization experiments). Images were deconvolved by SoftWoRx software using standard settings and the number of cells with Rad51 or Rfa1 foci or aggregates counted. Aggregates were identified as being at least 6 pixels (0.39 μm) wide, using ImageJ software.

#### Spindle pole bodies

To reduce disruption of the DAPI signal during the spreading process, cells were formaldehyde-fixed before spheroplasting. 0.5 ml 37% (v/v) formaldehyde was added to 4.5 ml of meiotic culture, and cells were incubated at room temperature for 30 min. Cells were harvested by centrifugation, washed with 5 ml 1% (w/v) potassium acetate, and were then processed as above for recombination protein detection through the PhotoFlo/water wash steps. Cover slips were affixed using Vectashield mounting medium with DAPI (Vector Laboratories), sealed with clear varnish and imaged on a DeltaVision microscope, (z=12-24, Exposure times: FITC=1.0 s, RD-TP-RE=1.0 s, DAPI=0.05-1.0 s). Images were deconvolved by SoftWoRx software using standard settings and the number of RFP (SPB) foci and DAPI signals per cell were counted, using GFP-tubulin as an additional signal where necessary.

### Recombination intermediate and product analysis

DNA was extracted, displayed on Southern blots, and hybridized with radioactive probe as described, either without (Oh et al., 2009); “Lichten lab” protocol) or with prior psoralen crosslinking (Kaur et al., 2018). JMs were detected using *Xmn*I digests probed with +156 to +1413 of ARG4 coding sequences. DSBs, NCOs, and COs were detected using *Eco*RI-*Xho*I digests probed with +539 to +719 of HIS4 coding sequences. Radioactive signal on blots was detected using either a Fuji LAS-3000 or a GE Typhoon FLA-9500 phosphorimager, and was quantified using Fuji Image Guage v4.22 software.

### Protein extraction and western blots

Samples containing 5-10 OD_600_ of cells were harvested by centrifugation, washed in 1 ml water, and flash frozen in liquid nitrogen. The pellet was stored at −80°C for at least 2 h, and was then resuspended in 150 μl 1.85 M NaOH, 7.5% (v/v) β-mercaptoethanol, and incubated on ice for 15 min. 150 μl of 55% Trichloroacetic acid was added and cells were incubated for a further 10 min. Cells were harvested by centrifugation for 10 min at top speed in a microfuge, and resuspended in 250 μl of 200 mM Tris-HCl pH6.5, 8 M urea, 5% (w/v) SDS, 1 mM EDTA, 0.02% bromophenol blue, 5% (v/v) β-mercaptoethanol, with 10 μl 25x protease inhibitor stock (Roche, 04 693 132 001). If necessary, 10 μl 1.5M Tris HCl pH8.8 was added to maintain pH, as shown by the blue indicator in the suspension. Cells were heat-shocked at 65°C for 10 min, centrifuged for 1 min at top speed in a microfuge, and the supernatant was harvested. Protein samples were denatured at 95°C for 5 min in appropriate loading buffer and loaded into precast SDS-PAGE gels (10% polyacrylamide; Biorad). Gels were run at 40 mA for 45 min in 1x running buffer (Fisher). Gels were assembled into a sandwich with Hybond C nitrocellulose membrane (Amersham) and Whatman 3MM blotting paper, and gel contents were transferred by semi-dry transfer (Thermo Scientific Pierce Power Blotter) using Pierce 1-Step Transfer Buffer for 7 min, or by wet transfer in 0.025 M Tris Base, 0.15 M Glycine, 2% (v/v) methanol at 150 mA for 2 h, or at 16V overnight (Biorad MiniProtean Tetra Cell). Membranes were rinsed in water. If necessary, the membrane was incubated 30 sec in Ponceau stain (Sigma), and then washed in PBST (Sigma). Membranes were incubated in 5% (w/v) skimmed milk in PBS at 4°C for 2-20 h, then incubated in primary antibody in 1% (w/v) skimmed milk in PBS at the following concentrations for 2-20 h at 4°C (anti-Srs2, Santa Cruz sc-1191, 1:2000; anti PSTAIR, Sigma P6962, 1:2500). Membranes were washed 3x in PBS, 5 min each at RT, and incubated in horseradish peroxidase-conjugated secondary antibody in 1% (w/v) skimmed milk in PBS at the following concentrations for 30-120 min at RT (anti-goat, Santa Cruz sc-2020, 1:2500; anti-mouse, Santa Cruz sc-2005, 1:2500). Membranes were washed 3x in PBS, 5 min each at RT, incubated with 2 ml high sensititivity ECL detection solution (Millipore), and blots were visualized on a GeneGnome chemiluminsescence imaging system.

## Supporting information

Supplemental Table S1-yeast strains

## End Matter

The authors declare no conflict of interest.

## Acknowledgments

We thank Douglas Bishop, Monica Boselli, Bin Hu, Adele Marston, and Akira Shinohara for strains, reagents, discussion, and communication of results in advance of publication, and Matan Cohen for technical assistance in preparing DNA samples. Imaging was performed using the DeltaVision microscope at the Wolfson Light Microscopy Facility, University of Sheffield. This work was supported by BBSRC BB/K009346/1 to ASHG, and by the Intramural Research Program at the Center for Cancer Research, National Cancer Institute, National Institutes of Health.

---

**Supplementary Figure S1.**
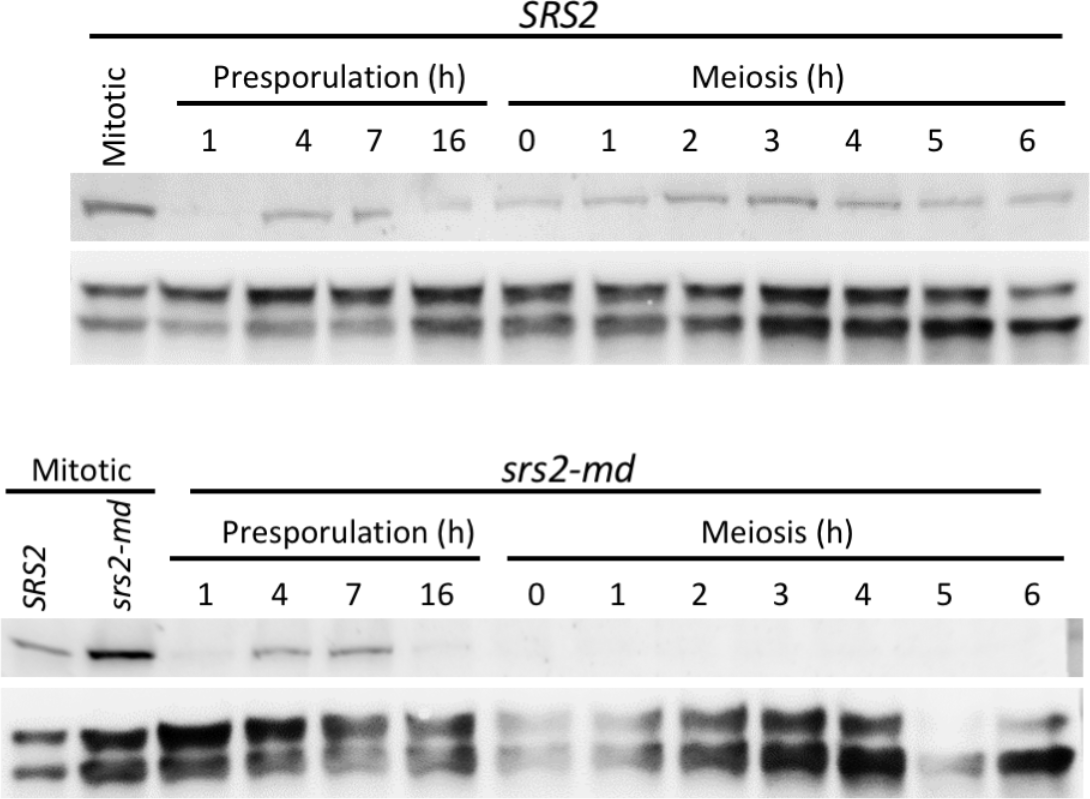
Srs2 loss during meiosis in *srs2-md* strains. Protein samples from mitotic (YPAD overnight), premeiotic (BYTA) or sporulation (SPM) cultures of *SRS2* or *srs2-md* strains were displayed on SDS-PAGE gels, transferred to nitrocellulose, and probe with anti-Srs2 or anti-PSTAIR as described in Materials and Methods. Cells were shifted to SPM after 17 h of presporulation growth in BYTA.

**Supplementary Figure S2.**
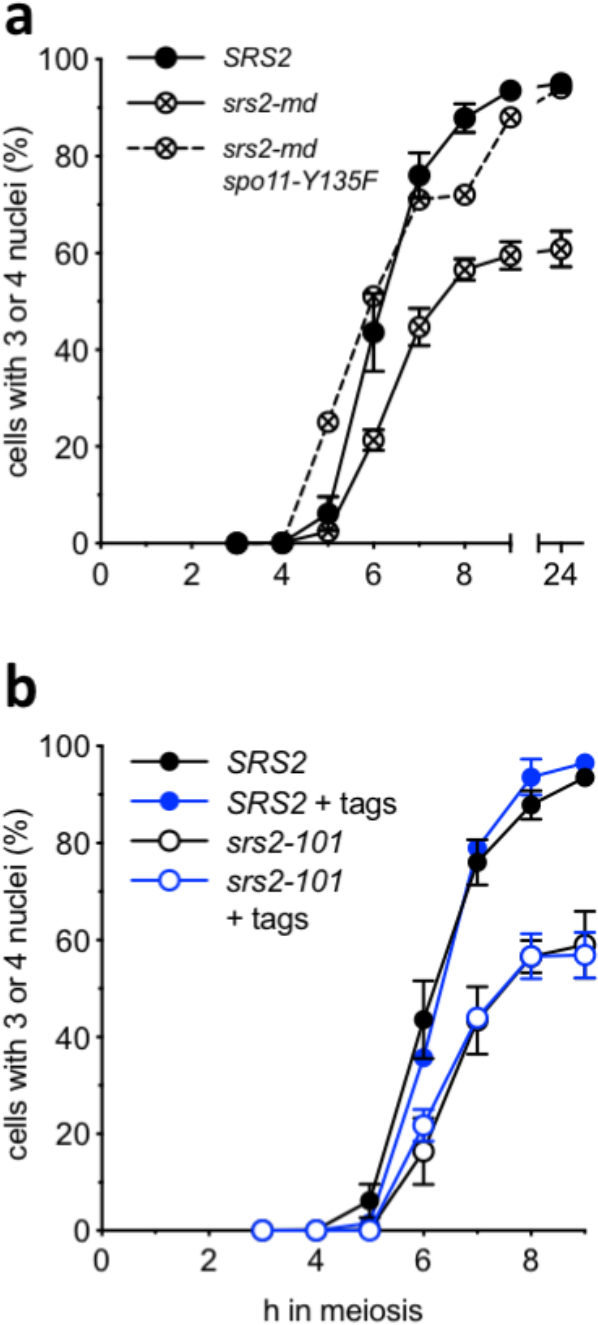
(a) Progression defects in *srs2* mutant cells require Spo11-induced double strand DNA breaks. While *srs2-md* mutants show a failure to complete meiosis II, *srs2-md spo11-Y135F* double mutants, which fail to form meiotic DSBs, restore progression to near wild-type levels (*srs2-md spo11-Y135F*, data from a single experiment;. data for *SRS2* and *srs2-md* reproduced from Figure 1). (b) Progression through meiotic nuclear divisions, expressed as percent of cells completing meiosis II, is similar in the presence (+ tags) or absence of *CNM67-mCherry* and *GFP-TUB1* (*SRS2* + tags, n=2; *srs2-101* + tags, n=2; error bars—range; data for untagged strains repeated from Figure 1).

**Supplementary Figure S3.**
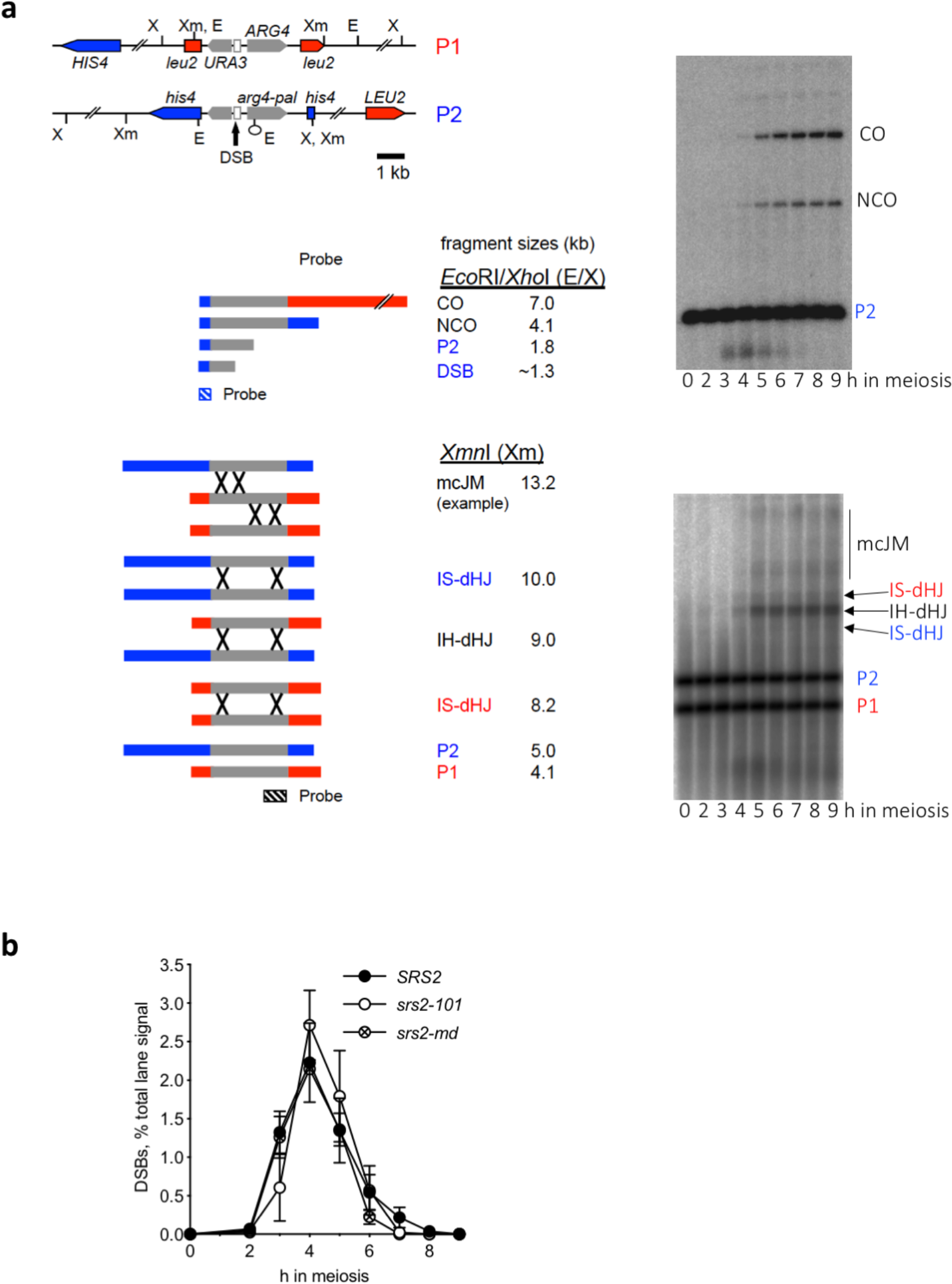
(a) Recombination reporter system used to detect recombination intermediates in *Xmn*I digests and products in *Eco*RI/*Xho*I double digests (Jessop et al., 2005; Jessop and Lichten, 2008). Grey arrows—*URA3* and *ARG4* genes; white box—63 nt telomere repeat sequence; vertical arrow—meiotic DSB hotspot. Inserts are at *LEU2* (red) on one chromosome III and at *HIS4* (blue) on the other. *arg4-pal* is an *Eco*RI-marked palindrome insertion for scoring noncrossovers. Relevant restriction sites are indicated. *Eco*RI/*Xho*I digests detect DSBs, NCO and CO products. X*mn*I digests detect interhomologue and intersister double Holliday junctions (IH-dHJs and IS-JMs, respectively), and multichromatid JMs (mcJMs, only one example of many possible is shown). Schematic reproduced from Kaur et al. (2015); illustrative Southern blots are from an *SRS2* diploid. (b) DSB dynamics are not altered in *srs2* mutants. DSBs were measured on Southern blots of *Eco*RI/*Xho*I digests of DNA from *SRS2* [n=5, data from De Muyt et al. (2012) and Kaur et al. (2015) and one additional replicate; error bars denote S.E.M.], *srs2-101* (n=5; error bars denote S.E.M.) and *srs2-md* (n=2; error bars denote range).

