## Supplemental Table S1-yeast strains for "*S. cerevisiae* Srs2 helicase ensures normal recombination intermediate metabolism during meiosis and prevents accumulation of Rad51 aggregates"

Supplementary Table S1. Yeast strains

| Strain Name | *MAT***a** Genotype | *MAT*$\alpha$ Genotype |
|---|---|---|
| dAG1668 | *ura3 ho::hisG leu2::hisG his4X arg4N srs2Δ::KanMX4* | *ho leu2 srs2Δ::KanMX4* |
| dAG1670 | *ho::LYS2 ura3 leu2 trp1::hisG his3::pHIS3-GFP-TUB1-HIS3 CNM67-3mCherry-NatMX4 srs2-101* | *ho::LYS2 ura3 leu2 trp1::hisG his3::pHIS3-GFP-TUB1-HIS3 CNM67-3mCherry-NatMX4 srs2-101* |
| dAG1681 | *ura3 lys2 ho::LYS2 leu2Δ(Xho1-Cla1) trp1::hisG srs2-101* | *ura3 lys2 ho::LYS2 leu2Δ(Xho1-Cla1) trp1::hisG srs2-101* |
| dAG1692 | *ho::LYS2 ura3 leu2 trp1::hisG his3:: pHIS3-GFP-TUB1-HIS3 CNM67-3mCherry-NatMX4* | *ho::LYS2 lys2 ura3 leu2::hisG his3-hisG trp1::hisG his3:: pHIS3-GFP-TUB1-HIS3 CNM67-3mCherry-NatMX4* |
| dAG1735 | *ho::LYS2 ura3 leu2 trp1::hisG his3::pHIS3-GFP-TUB1-HIS3 CNM67-3mCherry-NatMX4 srs2::KanMX* | *ho::LYS2 ura3 leu2 trp1::hisG his3::pHIS3-GFP-TUB1-HIS3 CNM67-3mCherry-NatMX4 srs2::KanMX* |
| dAG1756 | *ho::LYS2 ura3 leu2::hisG his3::hisG trp1::hisG* | *ho::LYS2 ura3 leu2::hisG his3::hisG trp1::hisG* |
| dAG1782 | *ho::LYS2 lys2 ura3 leu2 mek1::LEU2* | *ho::LYS2 lys2 ura3 leu2 mek1::LEU2* |
| dAG1783 | *ho::LYS2 lys2 ura3 leu2 mek1::LEU2 srs2-101::HphMX his4x* | *ho::LYS2 lys2 ura3 leu2 mek1::LEU2 srs2-101::HphMX* |
| dAG1813 | *ho::LYS2 lys2 ura3 leu2 pCLB2-3HA-SRS2::KanMX sae2::KanMX6 trp1::hisG arg4-nsp,bgl* | *ho::LYS2 lys2 ura3 leu2 pCLB2-3HA-SRS2::KanMX sae2::KanMX6 trp1::hisG his3::hisG* |
| dAG1814 | *ho::LYS2 lys2 ura3 leu2::hisG his3::hisG trp1::hisG pCLB2-3HA-SRS2::KanMX* | *ho::LYS2 lys2 ura3 leu2::hisG his3::hisG trp1::hisG pCLB2-3HA-SRS2::KANMX* |
| dAG1817 | *ho::LYS2 lys2 ura3 leu2::hisG trp1::hisG CNM67-3mCherry-NatMX4 his3::pHIS3-GFP-TUB1-HIS3 promURA3::tetR::GFP-LEU2-tetOx224-URA3* | *ho::LYS2 lys2 ura3 leu2::hisG trp1::hisG his3::pHIS3-GFP-TUB1-HIS3 CNM67-3mCherry-NatMX4* |
| dAG1818 | *ho::LYS2 lys2 ura3 leu2::hisG trp1::hisG CNM67-3mCherry-NatMX4 his3::pHIS3-GFP-TUB1-HIS3 pURA3::tetR::GFP-LEU2-tetOx224-URA3 pCLB2-3HA-SRS2::KANMX* | *ho::LYS2 lys2 ura3 leu2::hisG trp1::hisG his3::pHIS3-GFP-TUB1-HIS3 CNM67-3mCherry-NatMX4 pCLB2-3HA-SRS2::KANMX* |
| dAG1819 | *ho::LYS2 lys2 ura3 leu2::hisG his3::hisG trp1::hisG CNM67-3mCherry-NatMX4* | *ho::LYS2 lys2 ura3 leu2::hisG pURA3::tetR::GFP-LEU2-tetOx224-URA3 trp1::hisG CNM67-3mCherry-NatMX4* |
| dAG1820 | *ho::LYS2 lys2 ura3 leu2::hisG his3::hisG trp1::hisG CNM67-3mCherry-NatMX4 promURA3::tetR::GFP-LEU2-tetOx224-URA3 pCLB2-3HA-SRS2::KANMX* | *ho::LYS2 lys2 ura3 leu2::hisG his3::hisG trp1::hisG CNM67-3mCherry-NatMX4 pCLB2-3HA-SRS2::KANMX* |
| dAG1821 | *ho::LYS2 lys2 ura3 leu2 his3::hisG trp1::hisG arg4-nsp,bgl sae2::KanMX6* | *ho::LYS2 lys2 ura3 leu2 his3::hisG trp1::hisG arg4-nsp,bgl sae2::KanMX6* |

| Strain Name | *MAT***a** Genotype | *MAT*$\alpha$ Genotype |
|---|---|---|
| dAG1838 | *ho::LYS2 lys2 ura3 leu2::hisG pCLB2-3HA-SRS2::KanMX spo11-Y135F-3HA-His6::KanMX4* | *ho::LYS2 lys2 ura3 leu2::hisG his3::hisG trp1::hisG pCLB2-3HA-SRS2::KanMX spo11-Y135F-3HA-His6::KanMX4* |
| dAG1845 | *ho::LYS2 lys2 ura3 leu2::hisG his3::hisG trp1::hisG sae2Δ::HphMX* | *ho::LYS2 lys2 ura3 leu2::hisG his3::hisG trp1::hisG sae2Δ::HphMX* |
| dAG1846 | *ho::LYS2 lys2 ura3 leu2::hisG his3::hisG trp1::hisG sae2Δ::HphMX pCLB2-3HA-SRS2::KanMX* | *ho::LYS2 lys2 ura3 leu2::hisG his3::hisG trp1::hisG sae2Δ::HphMX pCLB2-3HA-SRS2::KanMX* |
| dAG1882 | *ho::LYS2 lys2 ura3 leu2::hisG his3::hisG trp1::hisG RFA1-GFP::KanMX* | *ho::LYS2 lys2 ura3 leu2::hisG his3::hisG trp1::hisG RFA1-GFP::KanMX* |
| dAG1883 | *ho::LYS2 lys2 ura3 leu2::hisG his3::hisG trp1::hisG RFA1-GFP::KanMX pCLB2-3HA-SRS2::KanMX* | *ho::LYS2 lys2 ura3 leu2::hisG his3::hisG trp1::hisG RFA1-GFP::KanMX pCLB2-3HA-SRS2::KanMX* |
| dAG1884 | *ho::LYS2 lys2 ura3 leu2::hisG his3::hisG trp1::hisG RFA1-GFP::KanMX pCLB2-3HA-SRS2::KANMX sae2Δ::HphMX* | *ho::LYS2 lys2 ura3 leu2::hisG his3::hisG trp1::hisG RFA1-GFP::KanMX pCLB2-3HA-SRS2::KANMX sae2Δ::HphMX* |
| dAG1887 | *ho::LYS2 lys2 ura3 leu2::hisG arg4Δ(eco47III-hpaI) trp1::hisG his3::hisG ndt80Δ(Eco47III-BseRI)::KanMX6 pCLB2-3HA-SRS2::KanMX* | *ho::LYS2 lys2 ura3 arg4Δ(eco47III-hpaI) leu2::hisG ndt80Δ(Eco47III-BseRI)::KanMX6 pCLB2-3HA-SRS2::KanMX* |
| dAG1892 | *ho::LYS2 lys2 ura3 leu2::hisG trp1::hisG his3::pHIS3-GFP-TUB1-HIS3 CNM67-3mCherry-NatMX4 pCLB2-3HA-SRS2::KanMX* | *ho::LYS2 lys2 ura3 leu2::hisG trp1::hisG his3::pHIS3-GFP-TUB1-HIS3 CNM67-3mCherry-NatMX4 pCLB2-3HA-SRS2::KanMX* |
| dAG1898 | *ho::LYS2 lys2 ura3 leu2::hisG trp1::hisG spo11-Y135F-3HA-His6::KanMX4 sae2Δ::HphMX* | *ho::LYS2 lys2 ura3 leu2::hisG spo11-Y135F-3HA-His6::KanMX4 sae2Δ::HphMX* |
| dAG1537 | *ura3 lys2 ho::LYS2 leu2Δ(Xho1-Cla1) trp1::hisG ZIP1-GFP srs2-101* | *ura3 lys2 ho::LYS2 leu2Δ(Xho1-Cla1) trp1::hisG ZIP1-GFP srs2-101* |
| dAG1534 | *ura3 lys2 ho::LYS2 leu2Δ(Xho1-Cla1) trp1::hisG ZIP1-GFP* | *ura3 lys2 ho::LYS2 leu2Δ(Xho1-Cla1) trp1::hisG ZIP1-GFP* |
| MJL2984 | *ura3Δ(hind3-sma1) lys2 ho::LYS2 cyh2-z arg4Δ(eco47III-hpa1) leu2-R his4::URA3-tel-arg4-ecPal9* | *ura3Δ(hind3-sma1) lys2 ho::LYS2 cyh2-z arg4Δ(eco47III-hpa1) leu2-R::URA3-tel-ARG4* |
| MJL3811 | *ura3Δ(hind3-sma1) lys2 ho::LYS2 arg4Δ(eco47III-hpa1) leu2-R his4::URA3rev-tel-arg4-ecPal9 pac1(62528)-Sph1 srs2-101* | *ura3Δ(hind3-sma1) lys2 ho::LYS2 arg4Δ(eco47III-hpa1) cyh2-z leu2-R::URA3rev-tel-ARG4 srs2-101* |
| MJL3553 | *ura3 lys2 ho::LYS2 arg4Δ(eco47III-hpa1) ndt80Δ(Eco47III-BseRI)::KanMX6 pCDC5-CDC5-pFA6a-HphMX4-pGAL1-CDC5 cyh2-z leu2-R his4::URA3-tel-arg4-ecPal9* | *ura3::pGPD1-GAL4(848).ER::URA3 lys2 ho::LYS2 arg4Δ(eco47III-hpa1) ndt80Δ(Eco47III-BseRI)::KanMX6 leu2-R::URA3-tel-ARG4* |
| MJL3638 | *ura3Δ(hind3-sma1) lys2 ho::LYS2 cyh2-z arg4Δ(eco47III-hpa1) trp1::hisG leu2-R his4::URA3-tel-arg4-ecPal99 srs2-101* | *ura3Δ(hind3-sma1) lys2 ho::LYS2 cyh2-z arg4Δ(eco47III-hpa1) trp1::hisG leu2-R::URA3-tel-ARG4 srs2-101 ndt80Δ(Eco47III-* |

| Strain Name | *MAT***a** Genotype | *MAT*$\alpha$ Genotype |
|---|---|---|
| | *ndt80Δ(Eco47III-BseRI)::KanMX6* | *BseRI)::KanMX6* |
| 3875 | *ura3 ho::LYS2 ura3Δ(hind3-sma1) arg4Δ(eco47III-hpa1) leu2-R::URA3-tel-ARG4 kanMX- pCLB2-SRS2* | *ura3 lys2 ho::LYS2 arg4Δ(eco47III-hpaI) cyh2-z leu2-R his4::URA3-tel-arg4-ecPal9 KanMX-pCLB2-SRS2* |
